# The Mag-Click-Capture-Release Technology for Selective Capture and Release of Hepatocyte-Derived Extracellular Vesicles as Biomarkers for Liver Disease

**DOI:** 10.1101/2024.10.25.620223

**Authors:** Richell Booijink, Anouk Mentink, Larissa Jansen, Sven Mentink, Bo van Rein, Lieke Geraets, Jorinde Scholten, Maureen Brusse, Siyu Fu, Andre Boonstra, Ruchi Bansal

## Abstract

Chronic liver diseases, such as liver cirrhosis and hepatocellular carcinoma (HCC), present major global health challenges, often diagnosed late. Circulating extracellular vesicles (EVs), which carry disease-specific biomolecular cargo, is emerging as an early diagnostic and prognostic biomarker for several diseases including cancer. However, current EV purification methods including ultracentrifugation and size exclusion chromatography present several limitations. Here, we present the Mag-Click-Capture-Release Technology for selective capture and release of EVs that combines <u>mag</u>netic beads, trans-cyclooctene (TCO) and tetrazine (Tz) <u>click</u> chemistry, immuno(antibody)-based <u>capture</u> and disulfide-driven <u>release</u> of EVs. Importantly, the Mag-Click-Capture-Release Technology is customizable, whereby using specific antibodies conjugated to TCO antibodies, different EV subtypes can be selectively captured and released for further analysis. With our Mag-Click-Capture-Release Technology, we successfully isolated hepatocyte-derived EVs from human serum with good recovery, high specificity and purity when compared with standard ultracentrifugation. Validation in serum samples obtained from cirrhosis and HCC patients with alcohol-associated liver disease evidenced an increasing trend in hepatocyte-EV levels correlating with disease severity, suggesting potential for early diagnosis and prognosis. In conclusion, we present here the Mag-Click-Capture-Release Technology, a customizable and efficient approach for selective isolation of organ-, cell-specific, and disease-relevant EVs from biological samples that can be subsequently released for downstream molecular EV analysis and EV-related functional assays.

## Introduction

Liver diseases pose a significant and increasing burden on the global healthcare system, contributing to approximately 2 million deaths each year, which accounts for 4% of all deaths globally [1, 2]. The majority of these deaths can be attributed to liver cirrhosis, the end stage of chronic liver disease (CLD) [3], and hepatocellular carcinoma (HCC), the most common form of primary liver cancer [4]. Owing to the increasing prevalence of metabolic disorders including obesity and type 2 diabetes and alcohol consumption, mortality and morbidity rates associated with liver diseases are rising exponentially [1]. Irrespective of the onset, cirrhosis develops following a long period of inflammation, during which healthy liver tissue is replaced by fibrotic scar tissue leading to portal hypertension [5]. Initially, liver cirrhosis can be asymptomatic, referred to as compensated cirrhosis. It can progress to decompensated cirrhosis, with complications such as jaundice, ascites, variceal bleeding or hepatic encephalopathy, resulting in frequent hospital admissions, impaired quality of life, and death in the absence of a liver transplant [5, 6]. Due to the long asymptomatic phase, liver cirrhosis – and HCC – are often diagnosed at late disease stages, which significantly impacts treatment and survival of patients with liver disease [4, 7]. Moreover, the scarcity in treatment options underscores the need for diagnostic and prognostic biomarkers to guide treatment decisions.

Current guidelines for diagnosing liver disease often rely on a combination of blood tests [alanine aminotransferase (ALT); aspartate aminotransferase (AST); alkaline phosphatase (ALP); gamma-glutamyl transferase; alpha fetoprotein (AFP)], and non-invasive imaging techniques (ultrasound; magnetic resonance imaging (MRI); and elastography) [8–10]. However, limitations of these tests include limited specificity and high variability [11–13]. Moreover, small lesions may be missed using current imaging techniques, which can show variable sensitivity, particularly in individuals with significant abdominal fat accumulation [14, 15].

To overcome these challenges, circulating extracellular vesicles (EVs) have emerged as promising biomarkers in a variety of diseases, including cancers [16, 17]. EVs are heterogeneous, membrane-bound structures released from cells and tissues that carry and transport diverse types of cargo (RNAs, proteins, lipids, microRNAs) to other cells [17]. Based on their size and biogenesis, they can be divided into three main subtypes: *i)* apoptotic bodies, ranging from 10000 – 5000 nm, *ii)* micro-vesicles, ranging from 100 – 1000 nm and *iii)* exosomes, ranging from 30 – 200 nm. Herein, because of their small size and abundance in tissues, exosomes are thought to play crucial roles in intercellular communication. Their specific biomolecular cargo reflects the pathophysiological state of their parental cell, which makes them promising disease biomarkers [18]. Indeed, genomic and proteomic analysis of isolated plasma EVs showed the potential of EVs as biomarkers for diseases as cancer and Alzheimer’s disease [19, 20]. Moreover, in mice, changes in EV composition and number were detected at earlier disease stages than visible tissue damage or other clinical and histological indicators [21]. Biomarker-based EV research requires the isolation of EVs from other constituents of body fluids, such as plasma or serum. The most common EV isolation methods are differential ultracentrifugation (UC), density gradient centrifugation (DGC), and size-exclusion chromatography (SEC). These techniques primarily focus on separation based on density (UC and DGC) or size (SEC) [22]. However, total blood EV is composed of vesicles released from all cells and tissues, where 80-90% of is platelet-derived, limiting the diagnostic value of analyzing total EVs, crucial information from disease-derived tissue EVs gets lost in the bulk [23]. In addition, these techniques are unable to purify EVs from other plasma or serum particles with similar sizes or densities, such as platelets and lipoproteins, which further interferes with EV analysis especially EV enumeration [24, 25]. Therefore, EV isolation techniques wherein EVs are isolated based on cell or tissue-specific markers are being explored [26–29]. For this, the specific binding interactions between an antibody and antigen is utilized to isolate EVs with specific antigens from complex systems. Herein, researchers have focused on isolating EVs with general EV protein markers (tumor susceptibility gene 101, TSG101, CD63, CD81, CD9), as well as specific surface protein adopted by EVs from their parental cells [30–33]. Unfortunately, the first line of standard immunoaffinity-based assays is suboptimal for clinical use. These assays have limited purity in isolating EVs from serum and plasma, due to high concentrations of proteins and lipids able to interfere with the immunoaffinity-based assays [34]. Moreover, the antigen–antibody mediated EV capture in these assays is based on a non-cleavable covalent linkage. This makes elution of the captured EVs challenging, and thereby also limiting the downstream analysis and EV-based functional studies.

Here, we present an efficient technology for the capture and release of cell (hepatocyte)-specific EVs, with high yield, purity, specificity and reliability. To accomplish this, we integrated immunoaffinity labeling using hepatocyte selective antibodies, <u>mag</u>netic beads, a highly reactive and biocompatible <u>click</u> chemistry reaction between the motifs tetrazine (Tz) and trans-cyclooctene (TCO) motifs, immuno(antibody)-based <u>capture</u>, and disulfide-driven <u>release</u> referred to as the Mag-Click-Capture-Release Technology. The Tz-TCO “click” is an irreversible but biocompatible chemical reaction with high reaction specificity [35], exceptionally fast kinetics and long-term aqueous stability, making it very suitable for biological applications [36, 37]. Moreover, Tz and TCO groups do not react or interfere with other functional groups in biological samples but exhibit high efficiency in conjugating with each other [38, 39]. Therefore, we envision a highly selective and fast conjugation of EVs to antibody-clicked magnetic beads, which can be enriched by applying a magnetic field. Subsequent release of EV by the cleavage of the disulfide linkage introduced on the magnetic beads [40], result in pure intact EVs suitable for downstream analysis and functional studies.

In this study, we demonstrate that the Mag-Click-Capture-Release Technology can enrich hepatocyte (HepG2)-derived EVs using the surface markers asialoglycoprotein receptor 1 (ASGPR1; hepatocyte specific [41]), and/or Epithelial cell adhesion molecule (EpCAM; epithelial cell-specific [42]). Furthermore, the Mag-Click-Capture-Release Technology was validated using artificial serum samples and compared with one of the commonly used EV isolation methods - ultracentrifugation. Finally, using the Mag-Click-Capture-Release Technology, we isolated and counted hepatocyte-EVs from serum samples obtained from patients with liver cirrhosis, early- and late-stage HCC versus (non-liver disease) control individuals.

## Results

### Design of the Mag-Click-Capture-Release Technology for the enrichment and analysis of hepatocyte-specific EVs

The Mag-Click-Capture-Release Technology is composed of two main elements: (A) a Tz-functionalized magnetic bead containing a stable disulfide bond [introduced through Ortho-Pyridyl disulfide (OPSS) linkage] and (B) a trans-cyclooctene (TCO)-conjugated antibody (Figure 1). Herein, the choice of antibody (BSA-free) for TCO conjugation can be easily adapted based on the desired EVs to be isolated determined by the EV surface marker expression. We conjugated TCO to the ASGPR1 antibody, which specifically targets hepatocyte surface receptor, asialoglycoprotein receptor (ASGPR1). When magnetic beads-Tz and TCO-ASGPR1 (step 1, Figure 1) are introduced into the serum, the Tz and TCO molecules will “click” together [39], resulting in the capture of ASGPR1-specific hepatocyte-derived EVs onto the magnetic beads (step 2, Figure 1). Through magnetic enrichment (step 3, Figure 1), the EVs captured on the beads are separated and washed to remove other constituents of the serum. Subsequently, the disulfide linkage introduced on the magnetic beads can be cleaved by exposure to a disulfide cleaving agent (i.e., dithiothreitol, DTT) (step 4, Figure 1) [40, 43]. This process releases the EVs from the beads, resulting in the enrichment of hepatocyte-specific EVs from the serum of patients with liver cirrhosis or HCC.

**Figure 1.**
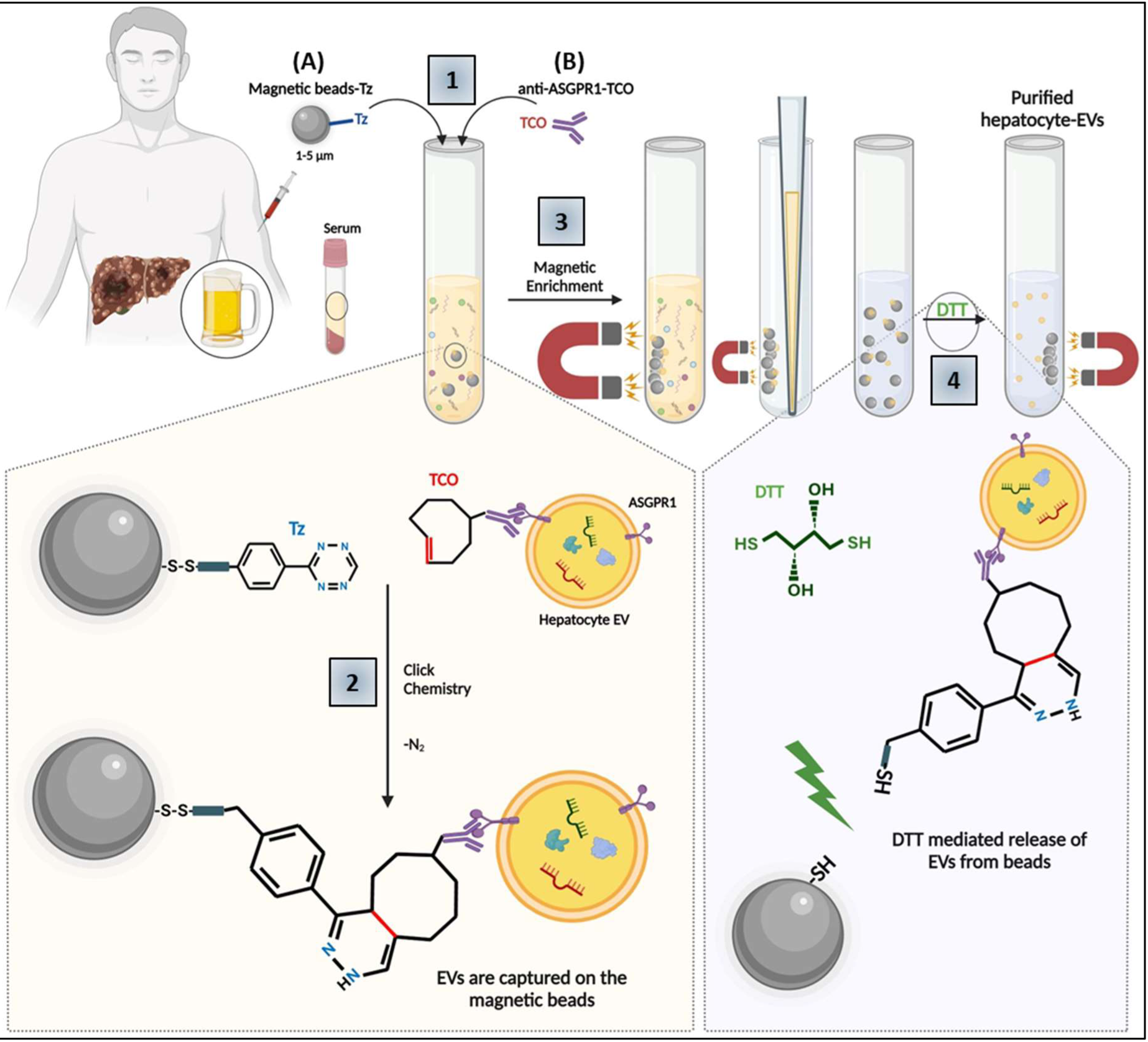
Purification of hepatocyte-specific extracellular vesicles using Mag-Click-Capture-Release Technology. 1) Magnetic beads (1-5 µm) functionalized with tetrazine (Tz) (A) and hepatocyte specific surface marker ASGPR1 conjugated to trans-cyclooctene (TCO) (B) are incubated into the serum of non-liver disease (healthy) individuals and (alcoholic) liver disease patients. 2) Under physiological conditions, Tz and TCO create a click reaction, resulting in the capturing of hepatocyte-EVs (bound to ASGPR1-TCO) to the magnetic beads, and 3) using a magnet, the hepatocyte-EVs can be enriched from other contaminants in the serum. 4) Purified hepatocyte-EVs can thereafter be released from the magnetic beads by hydrolyzing disulfide linkages using dithiothreitol (DTT).

### Optimization of the Mag-Click-Capture-Release Technology

All elements of the Mag-Click-Capture-Release Technology were optimized using flow cytometry. In this study, we used thiol-modified silica-coated magnetic beads (1-5 µm) with uniform size distribution. To synthesize Tz-conjugated magnetic beads, we first determined the optimal concentrations for OPSS (Figure 2A) and Tz (Figure 2B), by evaluating the mean fluorescence intensity (MFI) of the Bead-OPSS-Tz construct. The reactions (OPSS to thiol-modified magnetic bead and magnetic bead-OPSS to Tz, illustrated in **Supplementary Figure S1**) could be analyzed by Tz auto-fluorescence detectable in the orange/near-infrared spectrum as observed here and reported previously [44]. In both conjugations, the MFI reached a plateau around 3.8 mM as indicated by the dotted lines (Figure 2A-B), based on which, we adopted a concentration of 3.8 mM for OPSS and TZ as optimal for further conjugation reactions. Indeed, upon magnetic beads conjugation to OPSS followed by Tz, we observed a shift in the population of the magnetic beads in the fluorescent PE channel (excitation 488; emission 580/45) as indicated with an arrow (Figure 2C). The fluorescent images (in PE channel) were captured using high-resolution microscope which confirmed the successful conjugation of Tz on the beads (Figure 2C, insert). Finally, we evaluated the stability of Beads-OPSS-Tz and observed stable MFI for at least 15 weeks at 4 °C (Figure 2D) indicating that the Beads-Tz can be stored for at least 15 weeks at 4 °C without affecting its characteristics (Tz autofluorescence).

**Figure 2.**
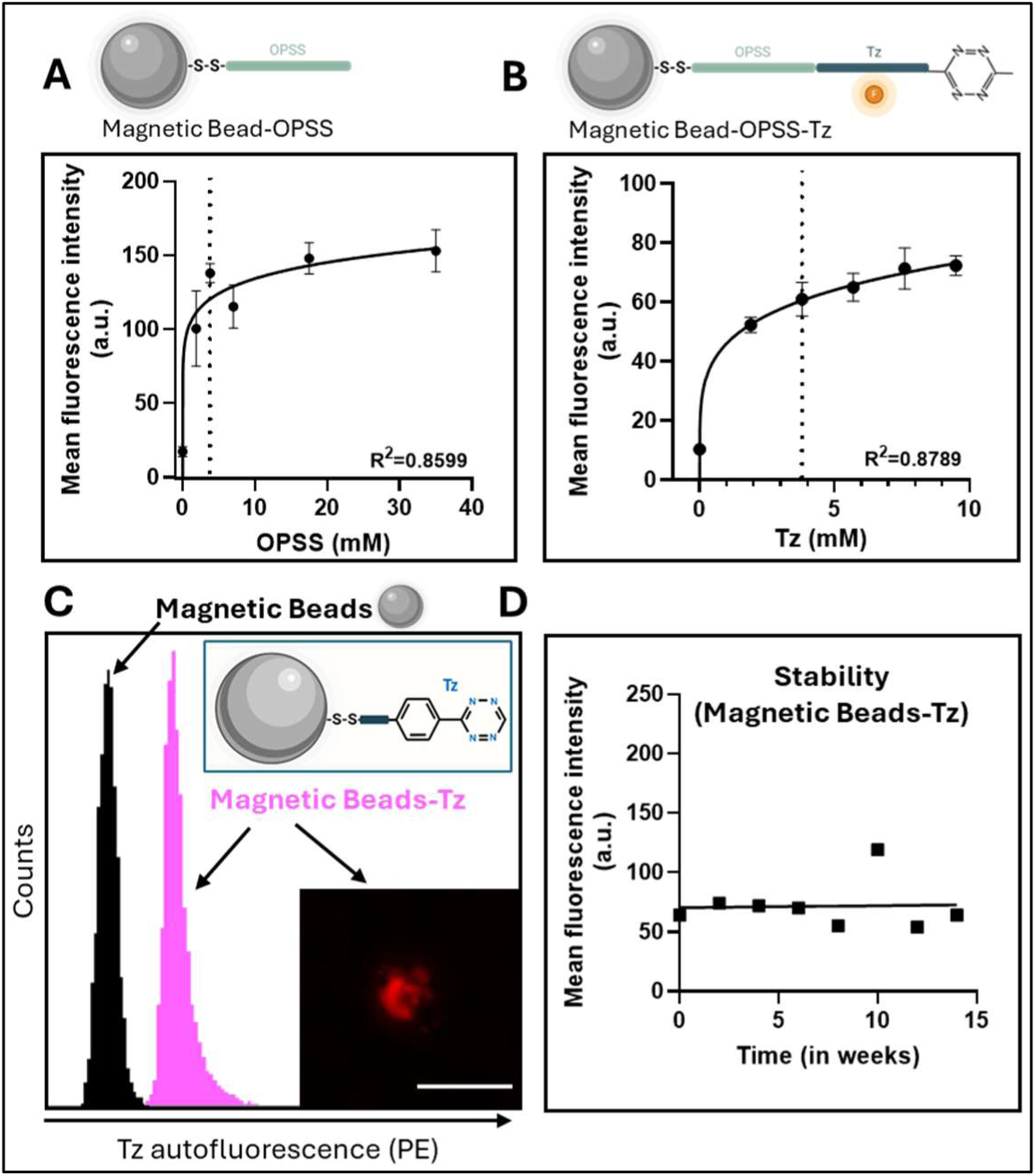
Optimization of the Mag-Click-Capture-Release Technology. A-B) Mean fluorescent intensity (MFI) of Tz (auto)fluorescence in the PE channel (488-585/40 nm) of Tz-OPSS-Beads with varying concentrations of A) OPSS and B) Tz, n=3. C) Representative flow cytometry histogram of empty magnetic beads (black) and beads-OPSS-Tz (purple), as well as fluorescent microscopy image, showing (auto)fluorescence of Tz-modified beads in the PE channel (488-585/40), n = 3, scale bar = 10 µm. D) MFI of Tz-OPSS-beads over time. Bars represent mean ± standard error of the mean (SEM).

### Optimization of the Mag-Click-Capture-Release Technology using EVs derived from HepG2

Successful capture of EVs using the Mag-Click-Capture-Release Technology was optimized using EVs derived from HepG2 (human hepatoma cell line) (Figure 3). To facilitate selective imaging of these nanometer-sized EVs, EVs were fluorescently labeled with Calcein-AM [45], a non-fluorescent dye that become fluorescent and impermeant when passively enters EVs [46], which allowed us to quantify the number of beads with fluorescent EVs attached to their surface using flow cytometry (Figure 3A). HepG2 EVs were isolated as per the method described in the methods section and illustrated in **Supplementary Figure S2.** HepG2 EVs were characterized for size and morphology using transmission electron microscopy (Figure 3B), Calcein staining by fluorescence microscopy (indicated by arrows, Figure 3C), and size distribution and concentration using NanoSight (Figure 3D). For the selective capture of HepG2 EVs, ASGPR1-TCO (Figure 3E) and EpCAM-TCO (Figure 3F) were freshly prepared (as detailed in the methods section, illustrated in **Supplementary Figure S1**) using two antibody concentrations (0.5 and 10 µg/ml). As seen in **Figure 3E-3F**, no fluorescence was observed without the addition of the antibody **(black)**, while a big shift in fluorescence was observed with 10 µg/ml ASGPR1-TCO (**Figure 3E, peach**) or 10 µg/ml EpCAM-TCO (**Figure 3F, cyan**) used to capture the EVs. In comparison, minimal fluorescence is observed with 20x less antibody of ASGPR1-TCO (**Figure 3F, purple**) or EpCAM-TCO (**Figure 3E, blue**), indicating that 10 µg/ml antibody is needed for capturing ASGPR1 or EpCAM-expressing EVs.

**Figure 3.**
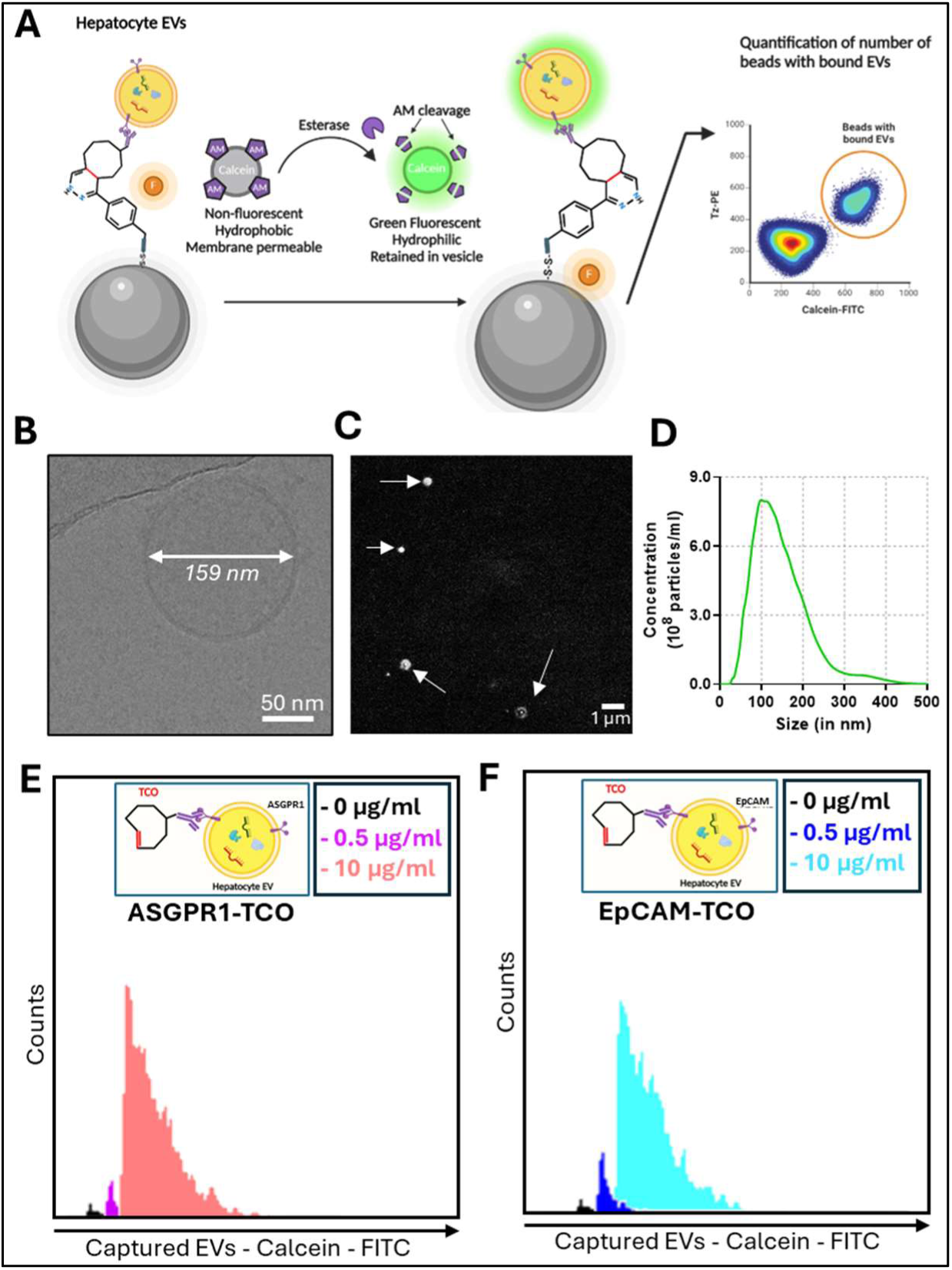
Optimization of Hepatocyte-EV Mag-Click-Capture-Release using HepG2-derived EVs. A) EVs captured on magnetic beads were fluorescently labeled with Calcein-AM, a non-fluorescent dye that become fluorescent by esterase and impermeant when passively enters EVs and can be quantified using FACS. B) Representative cryogenic transmission electron microscopy image of a HepG2 EV, scale bar = 50 nm. C) Representative fluorescent microscopy image of Calcein-AM-labeled HepG2 EVs, scale bar = 1 µm. D) Size-distribution of HepG2 EVs, analyzed by NanoSight, n=3. Representative flow cytometry histograms showing the number of EVs captured using 0, 0.5 and 10 µg/ml of ASGPR1 (E) and EpCAM (F), n=3.

### Enrichment of hepatocyte-EVs with the Mag-Click-Capture-Release Technology

Following the optimizations (**Figure 2** and **3**), we set out to determine the performance of the Mag-Click-Capture-Release Technology in an artificial setting where HepG2-derived EVs were introduced to PBS and normal human serum in a 1:10 ratio. For this, antibody-TCO and beads-Tz were added to Calcein-labeled hepatocyte EVs (**Figure 4A**, step 1). Successful capture of EVs to the beads was measured by flow cytometry (**Figure 4A**, step 2). Hereby, we counted the number of beads fluorescent for both PE (Tz autofluorescence) and FITC (Calcein-EVs), resulting in a percentage of beads that captured sufficient number of EVs to be detected by flow cytometry (**Figure 4A**). After enrichment, EVs were removed from the magnetic beads by the cleavage of disulfide linkage (**Figure 4A**, step 3), and the Calcein negative Tz-beads were measured again by FACS. We first isolated HepG2 EVs using ASGPR1 (hepatocyte-specific) (**Figure 4B**) and EpCAM (epithelial-specific) (**Figure 4C**) antibodies and saw a significant percentage of EVs were captured on the magnetic beads as depicted by flow plots (**Figure 4B-pink; 4C-blue**). This was further confirmed using fluorescent microscopy (**Figure 4D**) where Calcein-positive EVs were visualized in the FITC channel and beads-Tz were visualized in the PE channel (Tz) and in brightfield.

**Figure 4.**
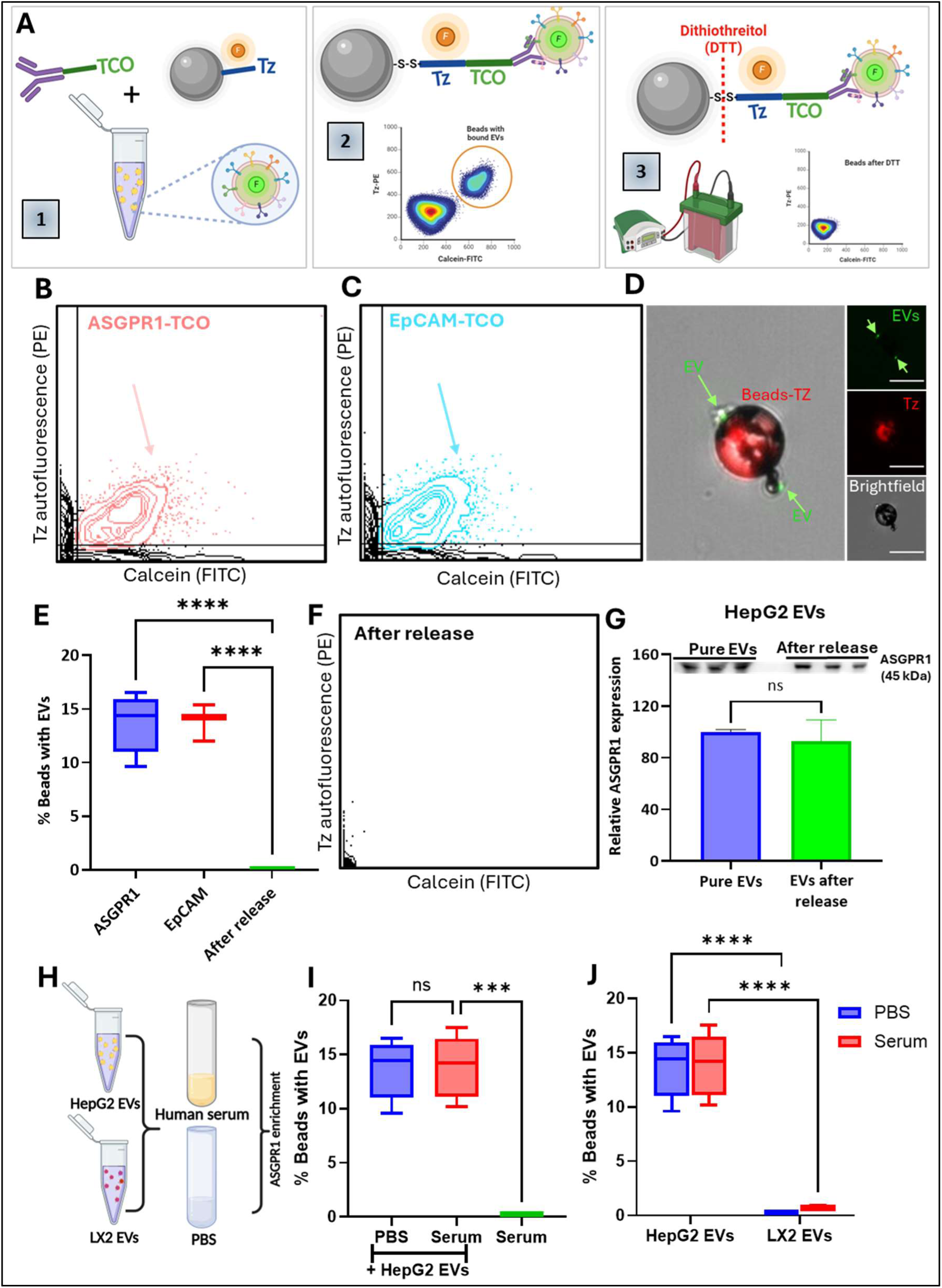
Enrichment of hepatocyte-EVs using the Mag-Click-Capture-Release Technology. A) Magnetic beads-Tz and antibody-TCO were added to fluorescently labeled EVs isolated from HepG2 cells (1), resulting in the capture of EVs on the magnetic beads (2) which were released from the beads by dithiothreitol (DTT) (2). The number of beads with EVs or without EVs (after release) were measured by flow cytometry (3), while the protein (ASGPR1) concentration in the released EVs is compared to the starting concentration of EVs by western blot (3). B-C) Representative scatter plots showing Tz autofluorescence versus Calcein positive EVs for B) ASGPR1 and C) EpCAM (blue) isolated EVs using the Mag-Click-Capture-Release Technology. D) Representative image of EVs captured on magnetic beads (Scale bar = 10 µm), showing the composite image, Calcein-stained EVs (green, indicated by arrows), Tz-fluorescent bead (red) and bead morphology (brightfield). E) Quantification of the percentage of beads with captured EVs on the surface, isolated by ASGPR1, EpCAM, and after DTT release, n=3. *F)* Representative scatter plot of the magnetic beads after DTT release, n=3. G) Relative ASGPR1 protein expression in HepG2 EVs before and after purification with the Mag-Click-Capture-Release Technology using ASGPR1, measured by Western Blot, n=3. H) HepG2 EVs or LX2 EVs were spiked in 1:10 ratio in human serum or in PBS followed by EV capture using Mag-Click-Capture-Release Technology. I) Percentage of beads with captured HepG2 EVs, isolated from PBS and Serum using ASGPR1 antibody, with and without spiking of HepG2 EVs. J) Percentage of beads with captured EVs from HepG2 cells or LX2 cells spiked in PBS or serum with the Mag-Click-Capture-Release Technology using ASGPR1. Bars represent mean ± standard error of the mean (SEM), n=3. Statistical analysis was performed by one-way or two-way ANOVA followed by a Tukey’s multiple comparisons test, and significant differences are denoted as *P < 0.05, ns denotes non-significant.

Using flow cytometry, we determined the percentage of beads with captured EVs, which was about 10-16% for both ASGPR1 and EpCAM (when only 10,000 beads were measured) (**Figure 4E**). Next, EVs were released from the beads using DTT, (**Figure 4A**, step 3) and the magnetic beads were measured using flow cytometry, which showed that the percentage of EVs attached to the beads dropped to near 0% (**Figure 4E-4F**).

Given that the goal of our research was to isolate hepatocyte-derived EVs, we proceeded with Mag-Click-Capture-Release EV enrichment using the hepatocyte-specific surface marker ASGPR1. To evaluate the yield of the Mag-Click-Capture-Release Technology, we measured ASGPR1 protein concentration of HepG2-EVs in PBS before and after enrichment and release, using Western Blot (**Figure 4A**, step 3). The relative ASGPR1 protein expression in enriched EVs was 93 ± 29 %, with respect to the ASGPR1 concentration in HepG2 EVs before enrichment, indicating high recovery of EVs following Mag-Click-Capture-Release EV enrichment and release (**Figure 4G**). A major challenge in isolating EVs from circulation is the abundance of interfering particles present in circulation and in biological samples e.g., serum that complicate antibody-based capture methods [34]. To assess the performance of our Mag-Click-Capture-Release Technology in circulation, we introduced HepG2-derived EVs into normal human serum at a 1:10 ratio (**Figure 4H**). Then, we compared EV capture in PBS (also diluted 1:10), artificial serum (containing HepG2-derived EVs), and control serum without the addition of HepG2-derived EVs (**Figure 4H**). Our results showed that the system has comparable capture efficiency in PBS and serum, while no EVs are captured in control serum without added EVs (**Figure 4I**). To further confirm the system’s specificity, we evaluated ASGPR1-mediated capture on EVs derived from human hepatic stellate cells (LX2 cells; non-hepatocytes purified using the same methods as HepG2). The Mag-Click-Capture-Release Technology showed nearly no capture of HSC-derived EVs in both PBS and serum (**Figure 4J**). These results demonstrate that, using the ASGPR1 antibody, our EV enrichment system is highly selective for hepatocyte-derived EVs.

### Performance of Mag-Click-Capture-Release Technology compared to ultracentrifugation

Currently, the gold standard for isolating EVs from serum or plasma is ultracentrifugation (UC) [22]. However, despite UC-based EV isolation is high throughput and applicable for many applications, the disadvantage is that it uses differences in density to isolate EVs from other contaminants. Therefore, UC-isolated EV samples contain the total EV population present in the sample, as well as all other particles with similar densities, such as lipoproteins, making analysis of a specific EV population challenging [47]. Here, we compared the UC-based EV isolation with our Mag-Click-Capture-Release Technology, in terms of specificity and purity of the analyte. To compare the specificity of the two techniques, we introduced HepG2 EVs into human serum, spilt the sample in half, and enriched one portion through ASGPR1-Mag-Click-Capture-Release while the other was enriched using UC (**Figure 5A**). Subsequently, total-EV were analyzed by Western Blot using EV marker TSG101 and hepatocyte-specific (ASGPR1) protein levels or by ELISA for ApoB lipoprotein contamination (**Figure 5A**). We observed similar ASGPR1 protein levels in UC and Mag-Click-Capture-Release enriched samples (**Figure 5B-C**). However, TSG101 protein levels were significantly higher in the UC-enriched samples (**Figure 5B-C**) suggesting that the amount of hepatocyte-EVs is similar in UC-enriched and Mag-Click-Release-enriched samples, while the total EV concentration is significantly higher in UC-enriched samples (**Figure 5C**). Consequently, the ratio of ASGPR1 (hepatocyte-EVs) compared to TSG101 (total EVs) is significantly higher in EVs isolated by Mag-Click-Capture-Release compared to EVs isolated by UC (**Figure 5D**), confirming that our Mag-Click-Capture-Release enrichment more selectively enriches hepatocyte EVs compared to UC. Next, we compared the purity of the Mag-Click-Capture-Release Technology with UC, by evaluating the concentration of Apolipoprotein B (ApoB) - a marker for (very-) low density lipoproteins (LDL/VLDLs) – in the samples [48]. For this, we isolated total EVs from human serum using UC or using Mag-Click-Capture-Release with the general EV marker CD63 conjugated to TCO and performed ApoB ELISA with the purified samples (**Figure 5A**). In the UC purified samples, the ApoB concentration was 500 ± 200 µg/ml, while in the Mag-Click-Capture-Release samples, the ApoB concentration was 60 ± 40 ng/ml (**Figure 5E**). This implies that in the Mag-Click-Capture-Release-purified samples, ApoB (VLDL/LDL) contamination is negligible, compared to UC-purified samples.

**Figure 5.**
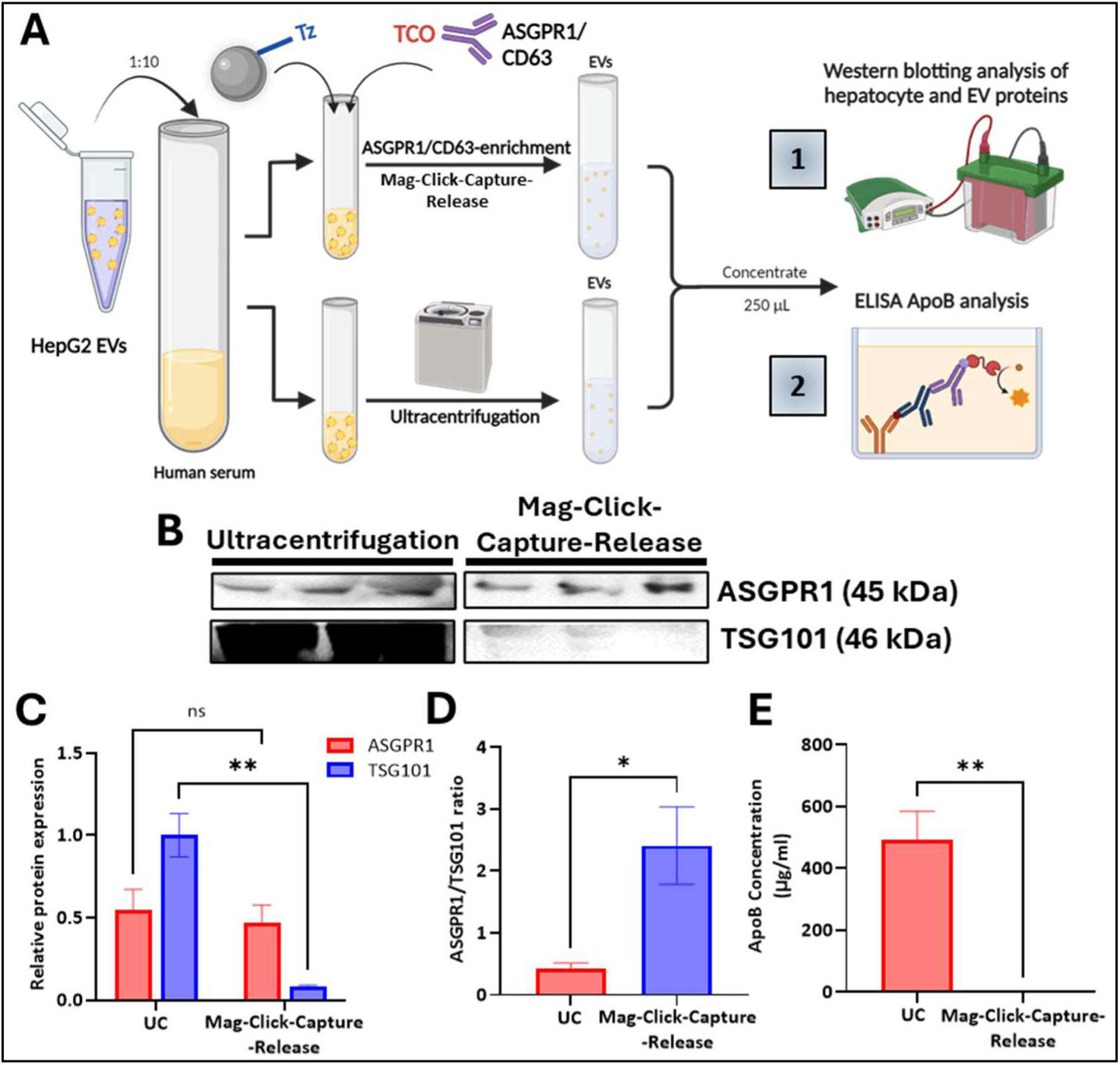
Comparison of the Mag-Click-Capture-Release Technology with ultracentrifugation. A) Schematic representation of the method used to compare the Mag-Click-Capture-Release Technology with ultracentrifugation. HepG2 cells were spiked into human serum in 1:10 ratio, and the sample was divided in two equal parts. In one part, EVs were purified using the Mag-Click-Capture-Release Technology with ASGPR1 antibody (Western Blot) or CD63 antibody (ELISA). In the other part, EVs were purified by ultracentrifugation (UC; 100.000 x g, 2h, 4 °C). Both purified samples were concentrated to equal volumes, and Western Blot was performed for ASGPR1 and TSG101 (1), or enzyme linked immunosorbent assay (ELISA) was performed for human apolipoprotein B (ApoB) (2). B-D) Western blot image (B) and quantification (C-D) showing ASGPR1, TSG101 and the ASGPR1/TSG101 ratio (D) in EVs purified with UC and the Mag-Click-Capture-Release Technology. E) Apolipoprotein B concentrations in EVs isolated by UC and CD63-Mag-Click-Release, measured by ELISA. Data is shown as mean ± SEM, n=3. Statistical analysis was performed using one way ANOVA followed by Tukey’s multiple comparisons test or an unpaired Student’s t-test. Significant differences are denoted as: ns = not significant, *P < 0.05, **P < 0.01.

### Purification of hepatocyte-EVs from liver cirrhosis and HCC patients

We collected 36 serum samples from ALD-related liver disease groups, including *(i)* early-stage cirrhotic HCC group, defined by Barcelona clinic liver cancer (BCLC [49]) stages 0-A, (*n* = 9; mean age = 66 y), *(ii)* late-stage cirrhosis HCC group, confirmed BCLC stages B-D, (*n* = 9; mean age = 72 y), *(iii)* cirrhosis group, (*n* = 9; mean age = 61 y), *(iv)* non-liver disease controls (*n* = 9; mean age = 42 y). The clinical characteristics of these cohorts are provided in **Supplementary Table S1**. For each clinical sample, 250 µl of aliquoted serum was introduced to the Mag-Click-Capture-Release Technology. The EVs captured on the beads were counted by FACS, and thereafter all ASGPR1-EVs were released from the samples. Upon increasing disease severity, we observed an increasing trend in the percentage of magnetic beads with captured hepatocyte-EVs (**Figure 6**).

**Figure 6:**
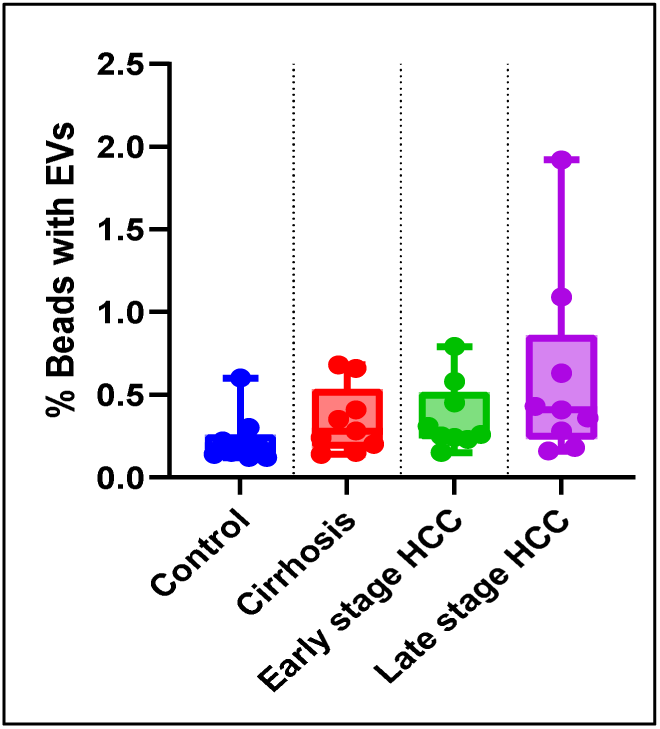
Isolation of Hepatocyte-EVs from liver cirrhosis and hepatocellular carcinoma patients. Percentage of beads (out of 10.000 beads measured) that captured hepatocyte-EVs from the serum of healthy donors (n=9), and patients with cirrhosis (n=9), early-stage hepatocellular carcinoma (n=9) and late-stage hepatocellular carcinoma (n=9). Data is presented as mean ± SEM.

Moreover, per 10.000 beads, the mean number of beads with EVs increased from 23 ± 15 in non-liver disease controls, to 61 ± 56 in late-stage HCC (**Table 1**), indicating an increased number of hepatocyte-derived EVs in the circulation of late-stage HCC patients. These results imply that upon disease progression, the number of hepatocyte-EVs in circulation increases.

**Table 1:** Number of beads with captured hepatocyte-EVs.

|  | Beads with EVs* |  |
| --- | --- | --- |
| | Mean ± SD | Median (range) |
| Healthy donors ($n = 9$) | $23 \pm 15.1$ | 19 (12 – 60) |
| Cirrhosis ($n = 9$) | $35 \pm 20.4$ | 28 (14 – 68) |
| Early cirrhotic HCC ($n = 9$) | $36 \pm 20.7$ | 26 (15 – 79) |
| Late cirrhotic HCC ($n = 9$) | $61 \pm 56.8$ | 41 (16 – 192) |

\* Per 10.000 beads

## Discussion

Circulating EVs have revolutionized the field of liquid biopsies, as they emerge as highly promising biomarkers due to their presence in biological fluids at early stages of disease, their inherent stability and the valuable cargo they carry from their parental cells. The selective isolation of EVs from biological fluids for biomarker discovery was previously investigated in the context of multiple diseases, including cancers [29, 31, 33, 50] and Alzheimer’s disease [51–53]. Moreover, liver-specific EV isolation from plasma using selective microbeads showed a significant trend in biomarker expression correlated to disease severity [32]. These studies suggest that EVs specifically enriched for their tissue of origin provide valuable insights into disease pathology and progression. However, the EV isolation techniques used in these studies are still suboptimal, as they are either based on advanced and costly microfluidic systems [29, 50], or immunoaffinity capture to beads or surfaces unable to release EVs [32, 33]. In this study, we have successfully developed and validated the Mag-Click-Capture-Release Technology. An easy-to-use, intuitive and customizable platform for the capture and release of specific EV subpopulations from human serum. This system combines the simplicity of magnetic bead-based purification, as previously done by others [32, 54, 55], with the high specificity of advanced covalent chemistry for EV capture and release [39]. This combination of EV magnetic bead-based capture and release, along with the ability to customize the selective antibody for each experiment, provides a great level of flexibility within the system which distinguishes the Mag-Click-Capture-Release Technology from current EV isolation methods [29, 32, 50].

By comparing the efficacy of the Mag-Click-Capture-Release in PBS versus particle-rich serum, as well as evaluating the non-specific binding of the Mag-Click-Capture-Release with a non-hepatocyte cell line, we demonstrated that the Mag-Click-Capture-Release Technology effectively purifies hepatocyte-EVs with high selectivity and specificity. 93% of the hepatocyte-EVs were recovered after release, verifying the high yield of the system. Moreover, we showed that our system outperforms conventional UC in the purification of a specific EV subpopulation from a biological sample such as human serum. Compared to UC, the Mag-Click-Capture-Release Technology isolates similar amounts of hepatocyte-EVs, confirming the EV yield is similar to UC, while the total number of EVs isolated is significantly reduced. Moreover, when we do indeed isolate all EVs using the Mag-Click-Capture-Release Technology with general EV-receptor CD63, lipoprotein contamination within the sample is reduced by 7000-fold. These findings again confirm the high selectivity and purity using our Mag-Click-Capture-Release Technology. This high selectivity and high yield are mostly attributed to the covalent chemistry-mediated click of the EVs to the beads, was previously shown to have superior binding properties compared to immunoaffinity-based capture [29]. This click chemistry reaction, an inverse-electron-demand Diels-Alder cycloaddition between Tz and TCO [35], is fast (rate constant of 10^4^ M^-1^ s^-1^ [56]), reactive, and stable without the addition of a catalyst. The Tz-TCO ligation is stable is specific, irreversible, and insensitive to biological influences such as enzymes, oxygen or water, resulting in the capture of EVs to the magnetic beads without the binding of nonspecific particles in the background. We initially demonstrated the versatility of the Mag-Click-Capture-Release Technology by successfully purifying similar amounts of EVs from the liver cancer cell line HepG2 using both an epithelial cell-specific antibody (EpCAM) and a hepatocyte-specific antibody (ASGPR1). Following this, we decided to further validate the Mag-Click-Capture-Release Technology for the purification of hepatocyte-derived EVs using ASGPR1. By selecting ASGPR1 as our capture surface marker, we intentionally focused on isolating hepatocyte-derived EVs from serum, rather than exclusively targeting epithelial-derived EpCAM-positive EVs [55]. ASGPR1, a transmembrane protein specifically expressed on hepatocytes, plays a crucial role in maintaining glycoprotein homeostasis [57]. However, in liver diseases, its role is under debate. For instance, one study found that ASGPR1 mRNA expression was upregulated in cirrhotic tissues, but significantly downregulated in HCC, correlating with increasing severity [58]. Conversely, ASGPR1 protein expression was upregulated in bulk EVs isolated from liver disease patients compared to healthy donors [59], and an increase in the number of ASGPR1 positive micro vesicles (EVs > 500 nm) in alcoholic hepatitis patients correlated with treatment response [60]. Similar to these studies, we observed an increasing trend in the number of isolated ASGPR1-EVs correlating with the disease severity. The differences within these studies could be due to different analytes examined. The first study analyzed ASGPR1 mRNA and protein levels in hepatic tissue [58], while others [59, 60] focused on ASGPR1 protein and receptor expression in circulating EVs. Thus, it has been hypothesized that while EVs can provide valuable information on the disease stage, they might have selective packaging, leading to variability in EV populations and possibly a discordance in protein and RNA levels between EVs and their parental tissue [61], which needs to be further evaluated.

We note that this study does have some limitations. By using flow cytometry as analytical method for counting the relative EVs captured, no exact EV count can be given. To do so, the number of EVs captured per bead needs to be evaluated, as well as the threshold for detecting fluorescence from the EVs. Unfortunately, accurately counting EVs, especially exosomes (100 – 200 nm), is difficult. Current methods including nanoparticle tracking analysis (NTA) or traditional flow cytometry instruments lack sensitivity, and are limited by a detection limit in particle concentration (NTA) or particle size (flow cytometry) [62]. Furthermore, while the EV counts in clinical samples showed an increasing trend related to disease severity, evaluation of the cargo of these EVs could provide additional, stronger, correlations between EVs and disease severity.

In future research, it may be useful to include a marker panel for the screening of liver cirrhosis and HCC. For this, micro-RNA’s (miRs) in EVs could be studied. For instance, in liver cirrhosis, an upregulation of miRNA-122 and miR-155 is observed in EVs [63], while miR-93, miR-224 and miR-665 are associated with HCC proliferation [64]. Moreover, EVs in ALD were found to carry high amounts of mitochondrial DNA (mtDNA), associated with triggering inflammatory signaling pathways and disease progression [65]. Moving forward, we aim to validate the Mag-Click-Capture-Release tool for diagnosing liver cirrhosis and HCC using larger study cohorts. Additionally, due to the system’s ability to efficiently release EVs while maintaining their cargo, we believe the Mag-Click-Capture-Release Technology could be employed to study EV-mediated communication networks, potentially between different tissues. For instance, labeled purified EVs could be introduced to other cells or tissues to examine their biodistribution and communication pathways under normal and pathological conditions.

To conclude, we successfully developed the Mag-Click-Capture-Release Technology, a customizable, easy-to-use and efficient system for the capture and release of specific EV populations from human serum. The system demonstrated the effective and selective isolation of hepatocyte-derived EVs with a yield of 93% and showed promise in correlating EV levels with disease severity in liver disease patients with ALD etiology. However, future research is needed to validate the Mag-Click-Capture-Release Technology as a tool for diagnosing liver cirrhosis and HCC. Lastly, its ability to customize the capture antibody and release EVs while preserving cargo could extend the use of the Mag-Click-Capture-Release Technology beyond biomarker studies for liver diseases.

## Acknowledgements

The authors would like to acknowledge all study participants (patients and their families, as well as healthy volunteers) for their contributions to this study. The authors would also like to thank the staff of the Personalized Diagnostics and Therapeutics (PDT) laboratory at the University of Twente, and the staff of the Department of Gastroenterology and Hepatology at Erasmus MC, for their contributions to this study. This work was supported by the University of Twente.

## Author contributions

RBo: Data curation; validation; investigation; visualization; methodology; writing – original draft; AM, SM, LJ, BvR, LG, and SF: Data curation; validation. AB: Writing – review and editing. RBa: Conceptualization; funding acquisition; writing – review and editing

## Competing interests

Authors declare no conflict of interest.

## Supplementary information

Supplemental information (Supplementary Figures S1 – S2; Supplementary Table S1) can be found online.

## STAR Methods

### 1. KEY RESOURCES TABLE

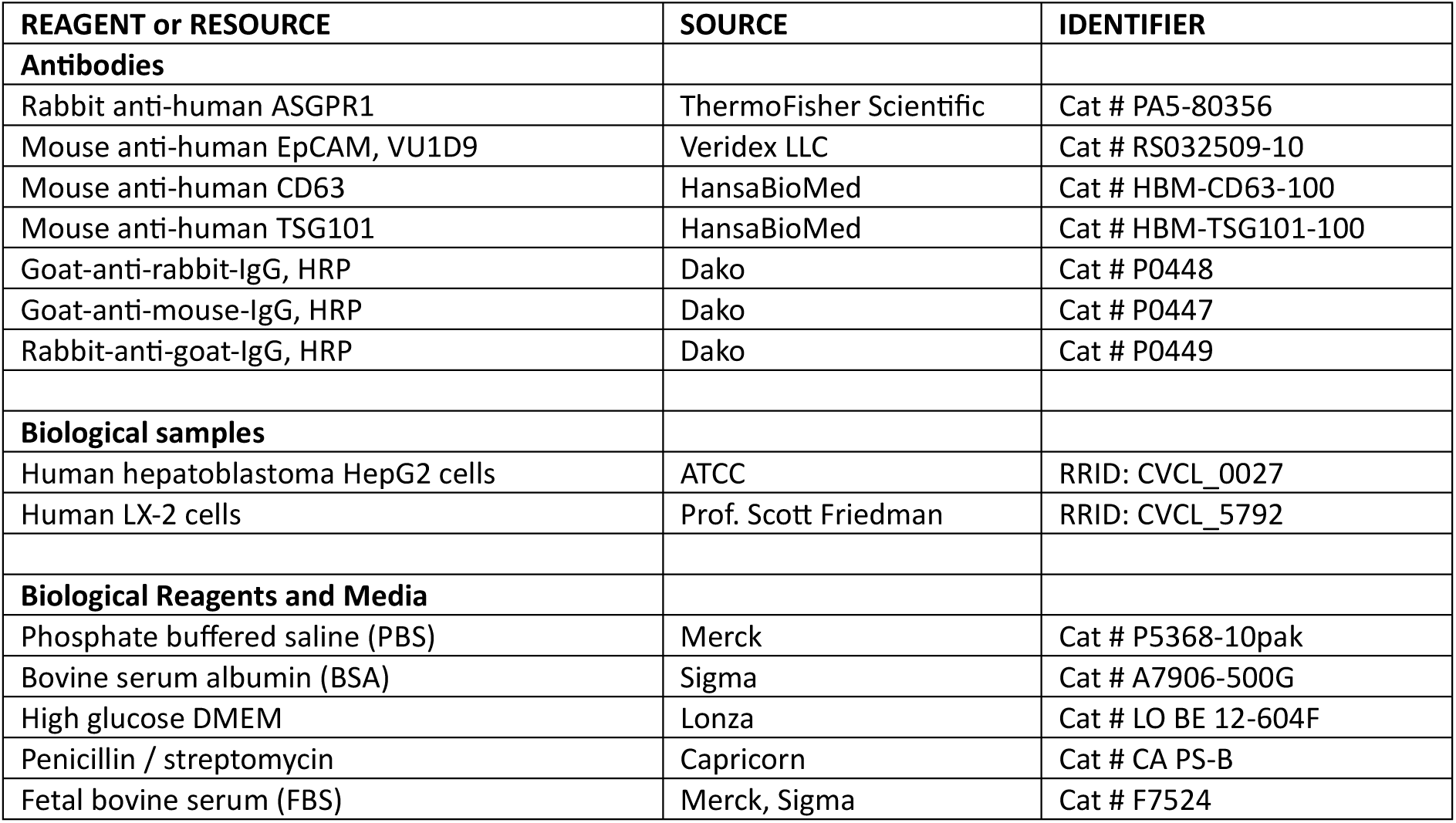
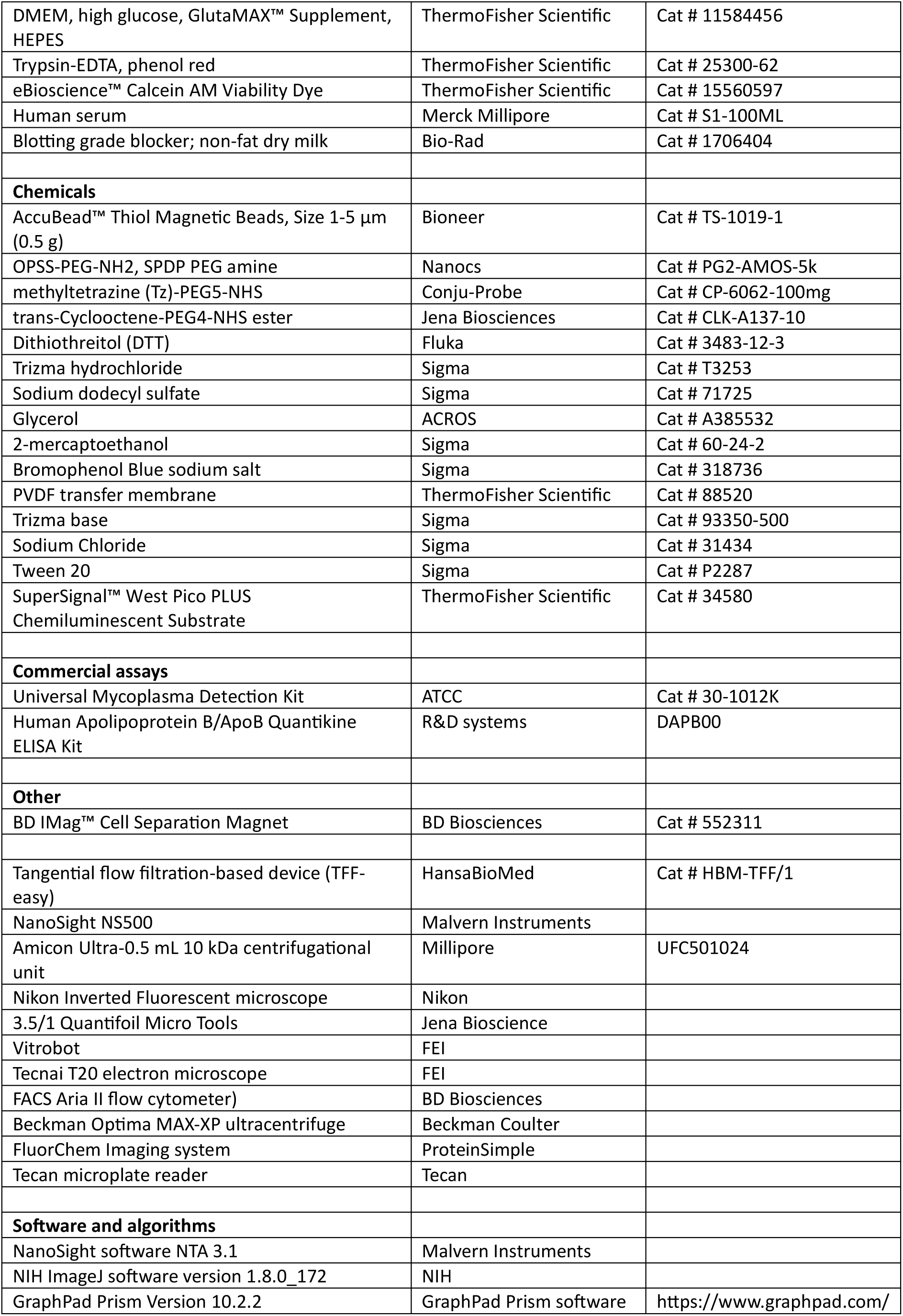

### 2. RESOURCE AVAILABILITY

#### Lead contact

Ruchi Bansal

#### Materials availability

The data is found in the figures and supplemental information in this paper. Moreover, the original flow plots are available as PDF upon request.

#### Data and code availability

Data: All data generated in the study is in the published article

Code: This data does not report original code.

The lead contact can provide information on all relevant data and recourses upon request.

### 3. STUDY PARTICIPANTS DETAILS

We collected a total of 36 serum samples from alcohol-related liver disease (ALD). The cohorts included the following groups: (i) early-stage cirrhotic HCC, defined by the Barcelona Clinic Liver Cancer (BCLC) staging system as stages 0-A (n = 9; mean age = 66 years), (ii) late-stage cirrhotic HCC, defined by BCLC stages B-D (n = 9; mean age = 72 years), (iii) cirrhosis without HCC (n = 9; mean age = 61 years), and (iv) non-liver disease control group (n = 9; mean age = 42 years). The clinical characteristics of each cohort were detailed in **Supplementary Table S1**. All clinical annotations were performed by a clinician who was blinded to the assay results to ensure objectivity.

### 4. METHOD DETAILS

#### Fabrication of Tz-conjugated magnetic beads and TCO-antibody conjugates – available upon request Cells

The human hepatoblastoma cell line HepG2 were cultured in high glucose containing Dulbecco’s modified Eagle Medium, supplemented with 10% Fetal Bovine serum (FBS) and antibiotics. The human hepatic stellate cell line LX-2 was provided by Prof. Scott Friedman. These cells were cultured in high glucose, GlutaMAX DMEM supplemented with 10% FBS and antibiotics. Cells were cultured at 37 °C, 5% CO_2_ and trypsinized when reaching 80-90% confluency using 0.05% trypsin-EDTA. All the cells were tested to exclude mycoplasma contamination using the Universal Mycoplasma Detection Kit thus ensuring that all experiments were performed with mycoplasma-free cells. Moreover, in our research, we adhere to strict laboratory practices to ensure the integrity and authenticity of our cell liens. We are confident about the reliability of our methods and the validity of the data produced from the cells used in this study.

#### Hepatocyte EV sample preparations

Cells, cultured to 90% confluency in a 175 cm^2^ culture flasks, were cultured in medium without FBS for 48 hours, to avoid any EV contamination from the serum added to the culture medium. Thereafter, the serum-free conditioned medium was collected and centrifuged at 300 x g for 10 minutes, followed by a second centrifugation at 2800 x g for 10 minutes. The supernatant was collected and concentrated to approximately 2 mL using tangential flow filtration, according to manufacturer’s instructions and visualized in **Supplementary Figure S2**. The concentrated EV sample was stored at 4 °C for short time use, or stored in aliquots at −80 °C.

#### Characterization of HepG2 EVs

To characterize the HepG2 EV, a T75 flask of HepG2 derived EVs was concentrated to 1,5 ml using TFF. Thereafter, the sample was loaded into the Nanosight, and analysis was performed according to the manufacturer’s instructions. Particle size distributions and concentration estimates were obtained using the accompanying software. To visualize the EVs, they were stained for 30 minutes at RT with 2 µM Calcein-AM viability dye and imaged by fluorescent microscopy. Moreover, cryogenic transmission electron microscopy (Cryo-TEM) was used to visualize the bilayers of the small EVs. Briefly, an aliquot (3 μL) of sample was deposited on 3.5/1 Quantifoil Micro Tools. After the excess liquid was blotted, the grids were vitrified in liquid ethane using a Vitrobot and transferred to a FEI Tecnai T20 electron microscope equipped with 20 a Gatan model 626 cryo-stage operating at 200 keV. Micrographs were recorded under low-dose conditions with a slow-scan CCD camera. The size of individual particles is quantified with NIH ImageJ software.

#### Magnetic-click enrichment of EVs

For optimization studies, HepG2- or LX2-EVs were fluorescently stained with 2 µM Calcein-AM for 30 minutes at 37 °C in the dark prior to enrichment (**Supplementary Figure S2**). Antibody-TCO conjugate (1:5 ratio) was added to the fluorescently labelled EVs in PBS/1% BSA buffer and incubated for 1 hour at room temperature on a shaking roller mixer. Thereafter, conjugated beads-Tz (1:10 ratio) were added and incubated for an additional hour on a shaking roller mixer. To remove unbound particles, the solution was washed three times with 5 minutes waiting steps using an IMag magnet, and the final Bead-EV conjugates were resuspended in PBS/1% BSA and samples analyzed by flow cytometry (FACS Aria II), after which EVs were released from the magnetic beads.

#### Magnetic-click enrichment of EV-enriched serum

EV-enriched serum samples were created by spiking concentrated HepG2- or LX2-EVs into human serum in a 1:10 ratio. EV-spiked human serum was incubated simultaneously with antibody-TCO (1:5 ratio) and Beads-Tz (1:10 ratio) overnight at room temperature on a shaking roller mixer. The following day, samples were washed 5 times with PBS/1% BSA with an IMag magnet, resuspended in 200 µL PBS 1% BSA and stained with 2µM Calcein-AM for 30 minutes at 37 °C. Samples were analyzed by flow cytometry and inverted microscopy, after which EVs were released from the magnetic beads.

#### EV release

To release the captured EVs from the magnetic beads, 50mM dithiothreitol (DTT), freshly prepared in PBS, was added in a 1:1 ratio to the coupled Beads-EVs and incubated for 30 minutes at room temperature on the vortex roller mixer. Thereupon, magnetic separation was used to separate the beads from the enriched EVs. The beads were run through FACS to confirm a successful EV release, and enriched EVs were stored in −80 °C until used for further characterization by Western Blot or Enzyme-Linked Immunosorbent Assay (ELISA) assays.

#### Fluorescent activated cell sorting

To evaluate successful conjugation of EVs to the magnetic beads, as well as quantify the number of beads with captured EVs, FACS was used. Hereby, 10,000 events were measured of each sample, where 1 bead is measured as 1 event. Conjugation of 6-methyl-tetrazine (Tz)-PEG_5_-NHS to the beads measured by the shift in fluorescent intensity in the PE (488 – 585/45 nm) channel. The percentage of beads successfully conjugated to EVs was estimated through analyzing the number of Calcein-AM positive events. Furthermore, the efficiency of EV-bead capture was observed by fluorescence microscopy (Nikon).

#### Ultracentrifugation

HepG2 EV-enriched or normal human serum was diluted 1:1 in particle free PBS. Samples were sequentially spun at 300 x g for 10 minutes, followed by 2800 x g for 10 minutes, and finally at 10,000 x g for an additional 10 minutes, with the supernatant collected after each step. Subsequently, samples were subjected to ultracentrifugation at 100,000 x g for 2 hours at 4 °C without break. Thereafter, the supernatant was discarded, and the pellet was resuspended in 1,5 ml PBS and stored in −80 °C. Prior to use, the samples were concentrated to 250 µL by centrifugation with centrifugal 10 kDa filters.

#### Western Blot

Samples (75 µL) were sonicated prior to mixing with 4x Laemmli loading buffer (1 M Tris-HCl pH 6,8; 2 % SDS, 40% Glycerol, 10% 2-mercaptoethanol, 0.05% Bromophenol Blue) and boiled at 95 °C for 5 minutes. Equal volumes of each sample (20 µL) were loaded only 12.5% SDS-PAGE gels and transferred to 0.2 µm polyvinylidene fluoride (PVDF) membranes. Nonspecific protein binding was blocked by 5% non-fat dry milk in Tris-buffered saline, 0.1% Tween 20 (TBST) for 1h at RT. Thereafter, membranes were incubated overnight at 4 °C with ASGPR1 or TSG101, diluted in blocking buffer. Blots were then incubated with horseradish peroxidase (HRP)-conjugated secondary antibodies for 1h at RT, followed by incubation with HRP-conjugated tertiary antibody for 1h at RT. Membranes were developed using SuperSignal™ West Pico PLUS Chemiluminescent Substrate and imaged by FluorChem Imaging System. The intensity of individual bands was quantified by ImageJ software.

#### Apolipoprotein B Enzyme-Linked Immunosorbent Assay

Total apolipoprotein B (ApoB) concentration was measured from 1 mL serum EVs isolated by UC or by Magnetic-Click enrichment with CD63 antibody using human ApoB ELISA Kit according to manufacturer’s instructions. Briefly, samples are incubated for 2h at RT on an ApoB pre-coated plate, followed by 2h incubation with human ApoB conjugate and 30 min incubation with substrate solution. The enzyme-substrate reaction was stopped by stop-solution, and optical density of is measured at 450 nm using a Tecan microplate reader. Sample concentrations were determined using a standard curve and adjusted for the dilution factors.

#### Clinical serum sample processing

TCO-conjugated anti-ASGPR1 (50 µl; 10µg/ml) and Beads-TZ (25 µl; ± 1,5 x 10^9^ beads) was added to 250 µL serum samples in 0.5 mL LoBind Protein tubes. The samples were incubated overnight at room temperature under continuous agitation. After magnetic enrichment, 20% of each sample was stained with 2µM Calcein-AM and analyzed with by FACS, to validate successful conjugation of EVs to beads and estimate the percentage of beads with EVs attached. The other 80% of each sample was released from the beads using 50mM DTT and stored at −80 °C for further analysis.

### 5. QUANTIFICATION AND STATISTICS

All data are presented as the mean ± standard error of the mean (SEM). Graphs and statistical analysis were performed using GraphPad Prism version 10.0.2 (GraphPad Prism Software, La Jolla, CA, USA). Comparisons between samples were analyzed using unpaired Student’s t-test or one-way analysis of variance (ANOVA) with Dunnett’s multiple comparisons. The differences were considered statistically significant as follows: *P < 0.05. **P < 0.01 and ***P <0.0001. All experiments were performed at least three times independently.

## Notes

### Competing Interest Statement

The authors have declared no competing interest.

