## Supplementary Figures and Table for "The Mag-Click-Capture-Release Technology for Selective Capture and Release of Hepatocyte-Derived Extracellular Vesicles as Biomarkers for Liver Disease"

Supplementary information

***Supplementary Figure S1: Schematic of preparing Tz-conjugated magnetic beads and TCO-conjugated antibodies.*** *Available upon request.*

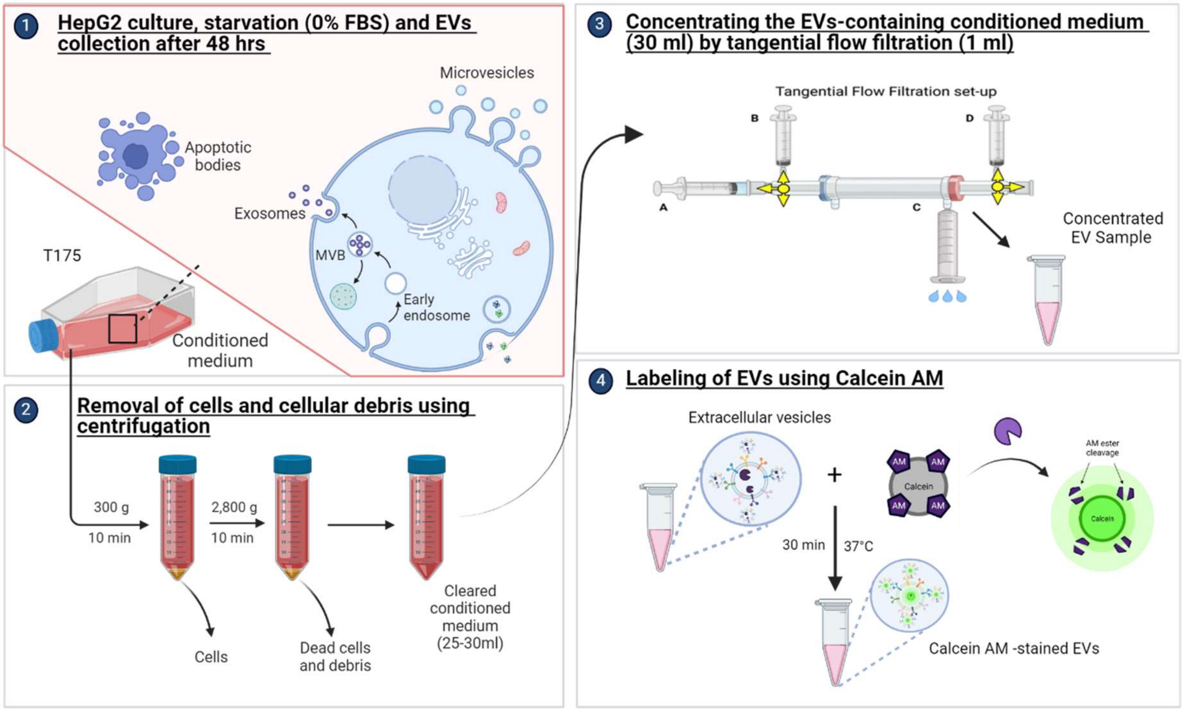


***Supplementary Figure S2: Preparation of HepG2-derived EVs.*** *1) In a T175 flask, HepG2 cells are cultured for 48h in starvation medium without FBS, and the conditioned medium is collected. 2) Cellular debris is removed by multiple centrifugation steps, after which the cleared conditioned medium is collected. 3) The conditioned medium is concentrated to 1 mL using tangential flow filtration (TFF). 4) HepG2 EVs are fluorescently labelled with Calcein-AM by incubation at 37 °C for 30 minutes.*

**Supplementary Table S1: Clinical characteristics cirrhosis and cirrhotic HCC patients.**

| | Cirrhosis (n=9) | Cirrhotic HCC | |
|---|---|---|---|
| | | Early (n=9) | Late (n=9) |
| **Sex** | | | |
| Male | 8 (89%) | 9 (100%) | 8 (89%) |
| Female | 1 (11%) | 0 | 1 (11%) |
| **Age** | | | |
| Median (range) - years | 66 (29 – 79) | 68 (48 – 77) | 72 (65 – 83) |
| **BCLC** | N.A. | | |
| 0 | | 2 (22%) | |
| A | | 7 (78%) | |
| B | | | 2 (22%) |
| C | | | 5 (56% |
| D | | | 2 (22%) |
| **Alpha fetoprotein, ng/mL** | | | |
| Mean ± SD | 4.8 (2.9) | 75 (207) | 180.53 (279.3) |
| Median (range) | 5 (1-10) | 6.8 (1 – 627) | 17 (4 – 588.1) |
| **Child Pugh score** | | | |
| Unknown | 3 | | |
| A | 3 | 6 | 6 |
| B | 3 | 2 | |
| C | | 1 | 3 |
